# Minor sequence modifications in temporin B cause drastic changes in antibacterial potency and selectivity by fundamentally altering membrane activity

**DOI:** 10.1101/312215

**Authors:** Giorgia Manzo, Philip M. Ferguson, V. Benjamin Gustilo, Tam T. Bui, Alex F. Drake, R. Andrew Atkinson, Giovanna Batoni, Christian D. Lorenz, David A. Phoenix, A. James Mason

## Abstract

Antimicrobial peptides (AMPs) are a potential source of new molecules to counter the increase in antimicrobial resistant infections but a better understanding of their properties is required to understand their native function and for effective translation as therapeutics. Details of the mechanism of their interaction with the bacterial plasma membrane are desired since damage or penetration of this structure is considered essential for AMP activity. Relatively modest modifications to AMP primary sequence can induce substantial changes in potency and/or spectrum of activity but, hitherto, have not been predicted to substantially alter the mechanism of interaction with the bacterial plasma membrane. Here we use a combination of molecular dynamics simulations, circular dichroism, solid-state NMR and patch clamp to investigate the extent to which temporin B and its analogues can be distinguished both *in vitro* and *in silico* on the basis of their interactions with model membranes. Enhancing the hydrophobicity of the N-terminus and cationicity of the C-terminus in temporin B improves its membrane activity and potency against both Gram-negative and Gram-positive bacteria. In contrast, enhancing the cationicity of the N-terminus abrogates its ability to trigger channel conductance and renders it ineffective against *Staphylococcus aureus* while nevertheless enhancing its potency against *Escherichia coli*. Our findings suggest even closely related AMPs may target the same bacterium with fundamentally differing mechanisms of action.

## Introduction

Antimicrobial peptides (AMPs) are a possible new source of antibiotic molecules and attract interest due to their multifaceted features which confer a broad spectrum of activity and low rate of resistance development.^1–3^ AMPs have been isolated from almost all living forms, from bacteria to humans, where they act as a primary line of defence against pathogenic infections.^4–6^ Often, a family of AMPs are found in each organism, sharing substantial primary sequence similarity and/or physical properties. It is generally accepted that the main factor determining the potency of AMPs is their interaction with the bacterial plasma membrane and that they kill pathogens by altering the permeation of the membrane and causing cell lysis^1–3,7^ or by crossing the membrane to interact with intracellular targets.^1,6,8^. Damage of model membranes by AMPs is well documented, with alteration of membrane permeability by AMPs explained using a variety of hypothetical models which have been added to and/or refined in recent years.^1,7^ It has been assumed that AMPs that are closely related will interact with bacterial plasma membranes with broadly similar mechanisms

The most common approach adopted to improve the antimicrobial properties of native AMPs, involves the modification of the physicochemical properties of the peptide sequence, including the net positive charge, hydrophobicity, hydrophobic moment and potential to adopt helical conformation to balance antimicrobial activity and host cell toxicity.^9^ Such studies often proceed on the basis that the overall mechanisms of bactericidal action are not expected to alter following minor sequence modification but that the native function of the AMP is enhanced. Recently we have compared the mechanism of action of seemingly closely related AMPs from the same class and found substantial differences in both the bacterial response to challenge and the ability of each peptide to penetrate model membranes.^10,11^ As a result, we hypothesised that even relatively minor alterations to the primary sequence of an AMP can result in fundamentally different behaviour when binding to models of the bacterial plasma membrane. The effects of an altered mechanism of action may be neutral but could lead to substantial changes in anti-bacterial potency and/or selectivity and may afford distinct native functions to even closely related AMPs from the same family.

To test our hypothesis, we report here our investigation of the effect of previously reported modifications to the sequence of temporin B on its ability to insert and disrupt model membranes and the effect this has on bactericidal activity against representatives of Gram-negative and Gram-positive pathogens. Temporin B (TB) belongs to the temporin family, one of the largest families of AMPs, whose members are closely related and were initially isolated from the skin mucus of the European common frog *Rana temporaria*.^12–14^ Temporins are mainly active against Gram-positive bacteria, with minimal inhibitory concentration (MIC) values ranging from 2.5-20 µM.^13,14^ They are less active against Gram-negative bacteria, with the exception of temporin L, which showed MIC ranging from 2-20 µM.^13,15^ Temporin B L1FK (TB_L1FK) and temporin B KKG6A (TB_KKG6A) are two analogues derived from the temporin B sequence, designed to improve activity against Gram-negative bacteria. Temporin B L1FK was obtained from a computational statistical model designed to predict the toxicity of the peptide as well as the antimicrobial activity. Temporin B L1FK maintains the physicochemical properties of the native peptide, such as hydrophobicity and amphipathic profile, but it has a broader spectrum of activity at low concentration, including against Gram-negative bacteria, as well as a higher potency in eradication of biofilms.^16,17^ Temporin B KKG6A was instead obtained by further optimization of an analogue obtained through alanine scanning of the temporin B sequence. The properties of temporin B KKG6A differ from temporin B, with a higher positive charge, a different amphipathic profile and a higher helical content, but is reported to have improved antimicrobial activity against Gram-negative and Gram-positive bacteria, a low haemolytic activity and a strong tendency to bind to lipopolysaccharide.^16,18^

To distinguish between the mode of interaction of the three temporin B analogues we combined both time-resolved and steady-state biophysical methods including all atom molecular dynamics simulations (using NMR structures of the three temporin B peptides solved in anionic detergent as a starting point), with circular dichroism and solid-state NMR to determine conformational and topological differences during the initial stages of membrane interaction and in the steady state. In addition, electrophysiology measurements of the current transitions on model membranes obtained with the patch-clamp technique reveal how these impact on the ability of each peptide to permeate analogous model membranes and enable understanding of the antibacterial outcome.

## Experimental procedures

### Materials

All the peptides (Table 1) were purchased from Pepceuticals (Enderby, UK) and Cambridge Research Biochemicals (Cleveland, UK) as desalted grade and were further purified using water/acetonitrile gradients using a Waters SymmetryPrep C8, 7 mm, 19 × 300 mm column. They were all amidated at the C-terminus. The lipids 1-palmitoyl-2-oleoyl-sn-glycero-3-phospho-(1′-rac-glycerol) (POPG), 1-palmitoyld_31_-2-oleoyl-sn-glycero-3-phospho-(1′-rac-glycerol) (POPG-d_31_), 1-palmitoyl-2-oleoyl-sn-glycero-3-phosphoethanolamine (POPE), 1-palmitoyld_31_-2-oleoyl-sn-glycero-3-phosphoethanolamine (POPE-d_31_), 1,2-diphytanoyl-sn-glycero-3-phospho-(1’-rac-glycerol) (DPhPG) and 1,2-diphytanoyl-sn-glycero-3-phosphoethanolamine (DPhPE) were purchased from Avanti Polar Lipids, Inc. (Alabaster, AL) and used without any purification. All other reagents were used as analytical grade or better.

**Table 1.**
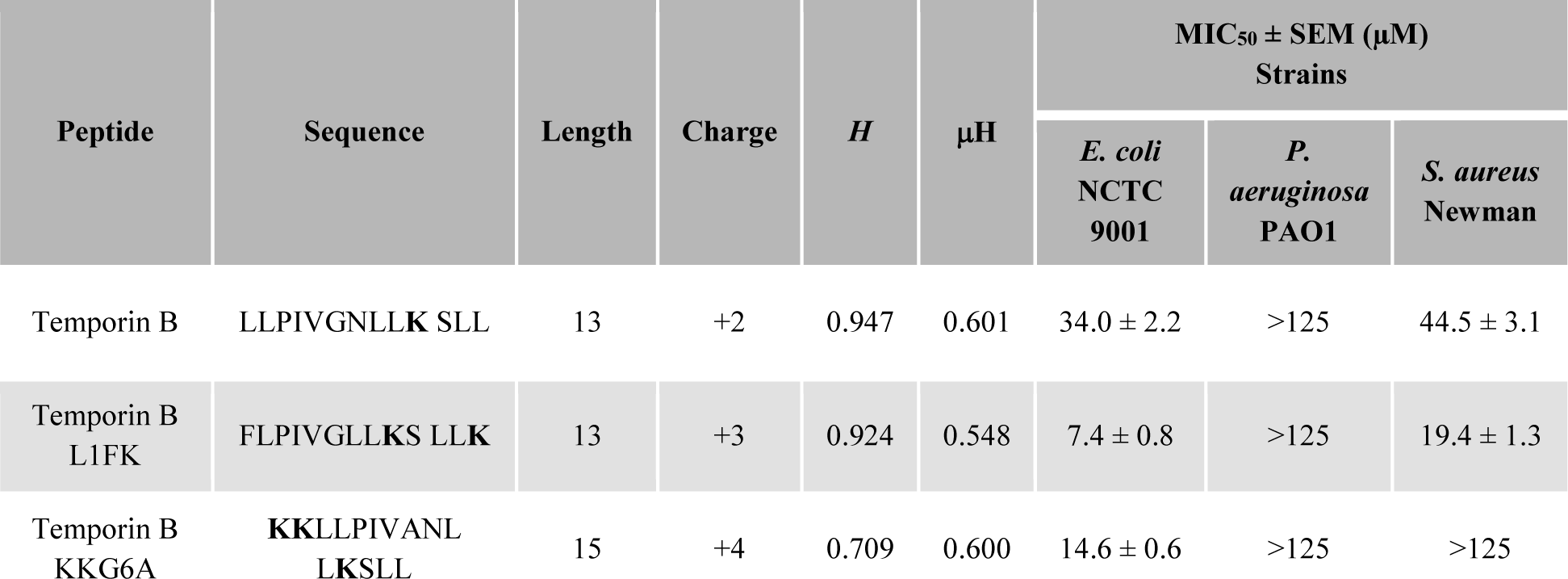
Peptides sequences, biophysical characteristics and antimicrobial activity against *Escherichia coli*, *Pseudomonas aeruginosa* and *Staphylococcus aureus*. All peptides were amidated at the C-terminus. Hydrophobicity (H) and hydrophobic moment (µH) were calculated using HeliQuest.^82^

### Antibacterial activity assay

Bacteria (*Escherichia coli* NCTC 9001, *Pseudomonas aeruginosa* PAO1 and *Staphylococcus aureus* Newman) were grown overnight in 5 mL of Mueller-Hinton (MH) broth without shaking at 37°C. The antibacterial activity of the five peptides studied was assessed through a modified two-fold broth microdilution assay, as described before.^10^ Briefly, a two-fold dilution of peptides’ stock solutions was performed in MH broth in a 96-well polystyrene microtiter (Greiner) plate. Then, 50 µL of a bacteria suspension (A_620_ of 0.0001) were added to 50 µL of peptide solution in each well of the plate reaching a final volume of 100 µL. Growth and sterility controls were present for each experiment. The plate was sealed and incubated at 37°C for 18 hours without shaking. Bacterial growth was observed with a colorimetric assay.^19^ 20 µL of a resazurin solution (0.2 g/L) were added to each well and the plate was incubated for further 2 hours. The results were read at A_570_ and A_600_ (pink and blue colour, respectively) and plotted as difference in absorbance versus peptide concentration. The plot obtained was fitted with a sigmoidal curve to obtain the concentration necessary to inhibit the growth of 50% of bacteria (MIC_50_). The MIC_50_ reported for each peptide in Table 1 is the average of at least three independent repeats.

### NMR spectroscopy and structure determination

Liquid state NMR spectroscopy experiments and data analysis were performed as described before.^11^ The samples solution consisted of 2 mM peptide and 100 mM deuterated sodium dodecyl sulphate (SDS-d_25_) micelles. 10% D_2_O containing trimethylsilyl propanoic acid (TSP) was added for the lock signal and as internal chemical shift reference. NMR spectra were acquired on a Bruker Avance 500 MHz spectrometer (Bruker, Coventry, UK) equipped with a cryoprobe. Standard Bruker TOCSY and NOESY pulse sequences were used, with water suppression using an excitation sculping sequence with gradients (mlevesgpph and noesyesgpph). The ^1^H 90-degree pulse length was 8.0s. One TOCSY with a mixing time of 60 ms, and two NOESY with mixing time of 100 and 200 ms were acquired for each sample. The relaxation delay was 1 s. 2048 data points were recorded in the direct dimension, and either 256 or 512 data points in the indirect dimension. CARA (version 1.9.1.2) and Dynamo software^20,21^ were used for the structure calculation. Inter-proton NOEs interactions were used as distance restraints in the structure calculation. CARA software generated a total of 200 structures on UNIO’08 (version 1.0.4)^22^ and XPLOR-NIH (version 2.40),^23,24^ after 7 iterations, using the software’s simulated annealing protocol. 20 structures with the lowest energy were chosen to produce a final average structure. In the case of ambiguous NOEs assignments by CARA software, Dynamo software’s annealing protocol was applied. Only unambiguous NOEs were used in this case after being classified as strong, medium and weak on the base of the relative intensity of the cross-peaks in the NOESY spectra. On the base of this classification an upper limit of 0.27, 0.33 and 0.50 nm have been applied, respectively, as restraint on the corresponding inter-proton distance, as described previously.^25^ One thousand structures were calculated and the 100 conformers with the lowest potential energy were selected for the analysis. The selected 100 conformers were aligned, and the root mean square deviation (RMSD) of the backbone heavy atoms was calculate with respect to their average structure. Solvent molecules were not included in the calculations. Structural coordinates were deposited in the Protein Data Bank (www.rcsb.org) under accession codes 6GIL, 6GIK and 6GIJ for temporin B, temporin B L1FK and temporin B KKG6A respectively.

### Molecular dynamic simulations

Simulations were carried out on either a Dell Precision quad core T3400 or T3500 workstation with a 1 kW Power supply (PSU) and two NVIDA PNY GeForce GTX570 or GTX580 graphics cards using Gromacs.^26^ The CHARMM36 all-atom force field was used in all simulations.^27,28^ The initial bilayer configuration was built using CHARMM-GUI.^29^ All membranes in this project contained a total of 512 lipids, composed either of POPE/POPG (75:25 mol:mol) or POPG to reflect the lipid charge ratios of the plasma membrane of Gram-negative and Gram-positive bacteria, respectively. Eight peptides were inserted at least 30 Angstrom above the lipid bilayer in a random position and orientation at least 20 Angstrom apart. The starting structures were taken from the NMR calculation in SDS micelles. The system was solvated with TIP3P water and Na+ ions added to neutralize. Energy minimization was carried out at 310 K with the Nose-Hoover thermostat using the steepest descent algorithm until the maximum force was less than 1000.0 kJ/ml/nm (~3000-4000 steps). Equilibration was carried out using the NVT ensemble for 100 ps and then the NPT ensemble for 1000 ps with position restraints on the peptides. Hydrogen bond angles were constrained with the Lincs algorithm. Final simulations were run in the NPT ensemble using 2 femtosecond intervals, with trajectories recorded every 2 picoseconds. All simulations were run for a total of 100 nanoseconds and repeated twice, with peptides inserted at different positions and orientations, giving a total of approximately 1.2 ms simulation.

### Liposomes preparation

Small unilamellar vesicles (SUVs) and multi-lamellar vesicles (MLVs) were prepared for circular dichroism (CD) and solid-state NMR (ssNMR) spectroscopy, respectively, and prepared as previously described.^11^ In both cases, lipids powders were solubilized in chloroform and dried under rotor-evaporation. To completely remove the organic solvent, the lipid films were left overnight under vacuum and hydrate in 5 mM Tris buffer with or without the addition of 100 mM NaCl (pH 7.0). Lipid suspension was subjected to 5 rapid freeze-thaw cycles for further sample homogenisation.

For ssNMR POPE/POPG-d31 (75:25, mol:mol), POPE/POPE-d31/POPG (50:25:25, mol:mol) and POPG/POPG-d31 (75:25,mol:mol) were used to prepare the vesicles. MLVs obtained were centrifuged at 21000 g for 30 minutes at room temperature and then pellets were transferred to Bruker 4 mm rotors for NMR measurements. Peptides were added to the buffer at a final concentration of 2% by mol, relative to lipids. POPE/POPG (75:25, mol:mol) and POPG SUVs were obtained by sonicating the lipid suspension on Soniprep 150 (Measuring and Scientific Equipment, London, UK) for 3 × 7 minutes with amplitude of 6 microns in the presence of ice to avoid lipid degradation. The SUVs were stored at 4C∘ and used within 5 days from preparation.

### Circular dichroism spectroscopy

Far-UV spectra of the peptides in the presence of SUVs and SDS micelles were acquired on a Chirascan Plus spectrometer (Applied Photophysics, Leatherhead, UK). Liposome samples were maintained at 310 K. Spectra were recorded from 260 to 190 nm. Lipid suspension was added to a 0.5 mm cuvette at a final concentration of 5.0 mM and then a few μl of a concentrated peptide solution were added and thoroughly mixed to give the indicated final peptide-to-lipid molar ratios. The same experimental conditions were used to investigate peptides secondary structure in SDS micelles. Final peptide concentration in the 0.5 mm cuvette was 40 µM, while SDS micelles concentration was 2 mM (L/P=50). In processing, a spectrum of the peptide free suspension was subtracted and Savitsky-Golay smoothing with a convolution width of 5 points applied.

### Solid state ^2^H-NMR

^2^H quadrupole echo experiments^30^ were performed at 61.46 MHz on a Bruker Avance 400 NMR spectrometer using a 4 mm MAS probe, spectral width of 100 KHz and with recycle delay, echo delay, acquisition time and 90° pulse lengths of 0.25 s, 100 μs, 2.6 ms and 3 μs respectively. The temperature was maintained at 310 K to keep the bilayers in their liquid-crystalline phase. During processing the first 10 points were removed in order to start Fourier-transformation at the beginning of the echo. Spectra were zero filled to 1 k points and 50 Hz exponential line-broadening was applied. Smoothed deuterium order parameter profiles were obtained from symmetrised and dePaked ^2^H-NMR powder spectra of POPG-d_31_ using published procedures (77, 78, 79).^31–33^

### Electrophysiology experiments (Patch-clamp)

Giant unilamellar vesicles (GUVs) composed of DPhPE/DPhPG (60:40, mol:mol) and DPhPG were prepared in the presence of 1 M sorbitol by electroformation method in an indium-tin oxide (ITO) coated glass chamber connected to the Nanion Vesicle Prep Pro setup (Nanion Technologies GmbH, Munich, Germany) using a 3-V peak-to-peak AC voltage at a frequency of 5 Hz for 120 and 140 minutes, respectively, at 36°C.^34–36^ Bilayers were formed by adding the GUVs solution to a buffer containing 250 mM KCl, 50 mM MgCl_2_ and 10 mM HEPES (pH 7.00) onto an aperture in a borosilicate chip (Port-a-Patch®; Nanion Technologies) and applying 70-90 mbar negative pressure resulting in a solvent-free membrane with a resistance in the GΩ range. After formation, a small amount of peptide stock solution (in water) was added to 50 µL of buffer solution in order to obtain its active concentration. All the experiments were carried on with a positive holding potential of 50 mV. The active concentration, the concentration at which the peptide first showed membrane activity, for each peptide was obtained through a titration performed in the same conditions. For all the experiments a minimum of 6 according repeats was done. Current traces were recorded at a sampling rate of 50 kHz using an EPC-10 amplifier from HEKA Elektronik (Lambrecht, Germany). The system was computer controlled by the PatchControl™ software (Nanion) and GePulse (Michael Pusch, Genoa, Italy, http://www.ge.cnr.it/ICB/conti_moran_pusch/programs-pusch/software-mik.htm). The data were filtered using the built-in Bessel filter of the EPC-10 at a cut-off frequency of 10 kHz. The experiments were performed at room temperature. Data analysis was performed with the pClamp 10 software package (Axon Instruments).

## Results

### Primary sequence and antibacterial activity

Comparison of the primary sequences of temporin B (TB) and its two analogues, temporin B L1FK (TB_L1FK) and temporin B KKG6A (TB_KKG6A) indicates the extent of the modifications to the primary sequence and their effects on nominal charge, hydrophobicity (H) and hydrophobic moment (µH), assuming each peptide adopts an idealised α-helix (Table 1). All three temporin B peptides are relatively hydrophobic and short and even single amino acid substitutions will be expected to have some impact on the physicochemical properties. Temporin B L1FK and temporin B KKG6A were both designed^16,18^ to broaden the spectrum of activity of temporin B. For temporin B L1FK, the optimization involved the substitution of Leu1 with Phe1 to render the N-terminus more hydrophobic, the addition of a lysine residue at the C-terminus and the deletion of Asn7.^16^ These modifications improved the activity of the peptide inducing a 4.5-fold increase in potency against uropathogenic *Escherichia coli* NCTC9001 and a two-fold increase against *Staphylococcus aureus* Newman. To create temporin B KKG6A, two lysines are added to the N-terminus, increasing the nominal positive charge and Gly6 is substituted with alanine.^18^ These modifications improved the activity of the peptide against *E. coli* NCTC9001 with slightly more than a two-fold improvement in potency, but the peptide loses activity against *S. aureus* Newman. None of the three temporin B peptides were effective against *Pseudomonas aeruginosa* PAO1 in the conditions used here. The results reported here for temporin B KKG6A are in disagreement with those reported previously, which showed up to 8-fold improvement in the activity against Gram-positive bacteria^16,18^ and 16-fold against *P. aeruginosa*^16^ when compared to temporin B. However different bacterial strains and protocols were applied in the previous work. Although limited in scope, the antibacterial susceptibility testing indicates that, indeed relatively small changes in the primary sequence can have substantial impacts on antimicrobial activity and, further, that these impacts cannot be easily related to changes in physicochemical properties such as nominal charge or hydrophobicity. Specific effects related to the peptide conformation or ability to effectively penetrate membranes can be suspected.

### Conformation and structure of temporin B peptides in sodium dodecyl sulphate (SDS) micelles and in model membranes in the steady state

As a first step, the secondary structures of the three temporin B peptides were initially analysed and characterised with far-UV CD and NMR spectroscopy in the presence of SDS micelles to mimic the negative surface of a bacterial membrane (Fig. 1). Structures for temporin B and temporin B KKG6A have been solved previously in LPS micelles^18,37^ and in the presence of *E. coli* and *Staphylococcus epidermidis* cells^38,39^ although coordinates are not available in the Protein Data Bank.

**Figure 1.**
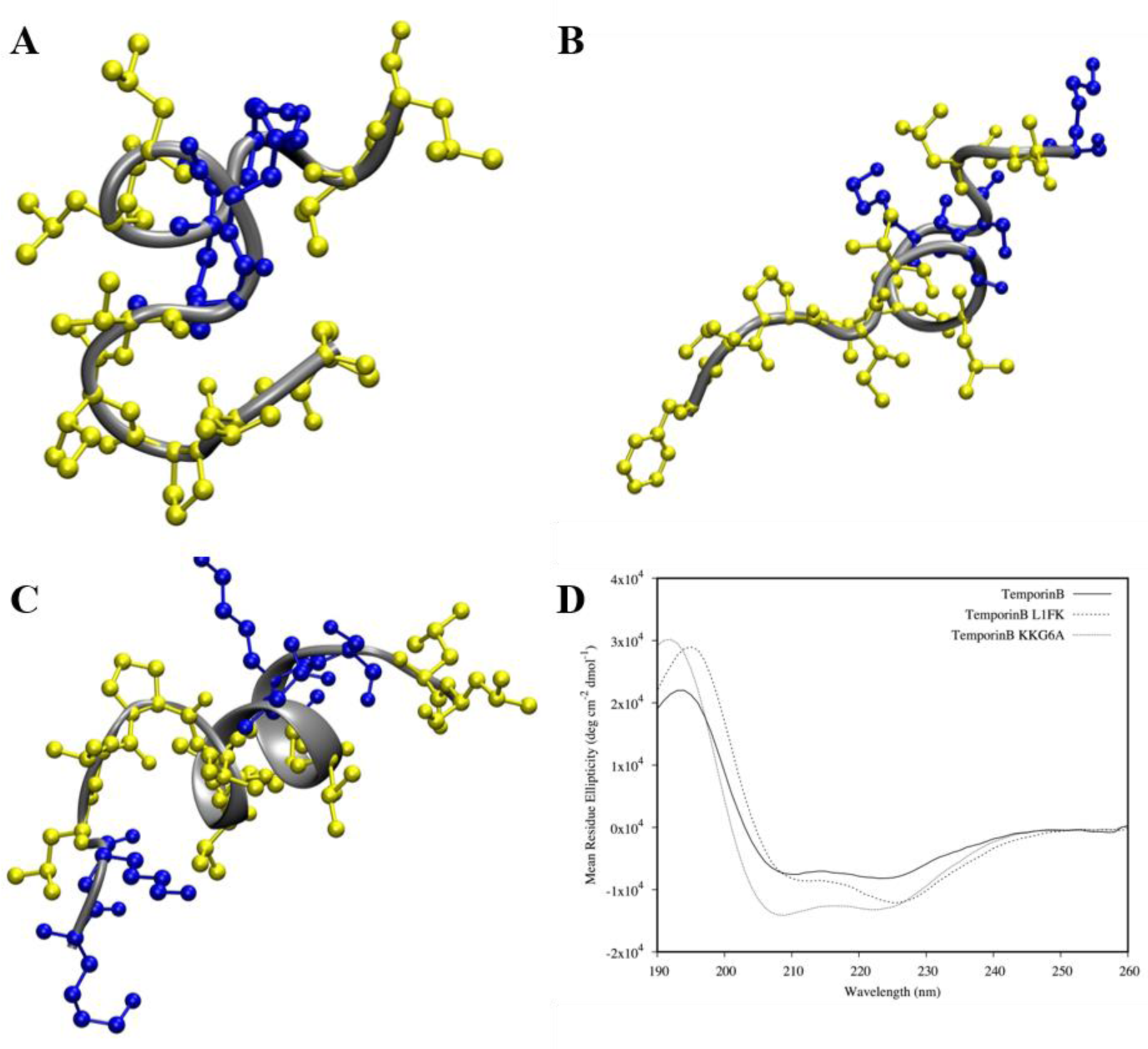
Three-dimensional structure for the three temporin B analogues. Structures were determined through ^1^H-NOESY NMR spectroscopy in SDS-d_25_ micelles (peptide/detergent equal to 1/50). The structure with the minimum RMSD is shown and was used as starting point in the MD simulations. The hydrophobic and hydrophilic residues are shown in yellow and blue respectively for temporin B (A), temporin B L1FK (B) and temporin B KKG6A (C). CD spectra of the AMPs in the same conditions used for NMR experiments are shown in (E). NMR and CD spectroscopy both revealed a lower propensity in helical folding for TB and TB L1FK peptides. NMR structures were used as starting configurations for MD simulations of model membranes.

Temporin B KKG6A adopts a more extended double β-turn (~40% of the whole sequence) in the -IVANLK-region. In all three peptides there is little evidence of order from the N-terminus to the glycine residue (temporin B and temporin B L1FK) or from the N-terminus to the proline residue (temporin B KK6A). In support, the far-UV CD spectra indicate that the Ramachandran angles adopted by the ordered regions of the peptides are similar to those of an α-helix and hence an α-helix like spectrum. The greater magnitude of the temporin B KKG6A CD spectrum compared with the temporin B CD spectrum reflects the increase from ~27% to ~40% whole molecule order. In contrast, the CD spectrum of temporin L1FK is more compatible with the extended α-helix often found in membrane proteins located in membranes as opposed to just water.^40^ These results are in broad agreement with the predicted effects of the design of temporin B L1FK^16^ where the aim was to largely retain the physicochemical properties of the ancestor peptide.

As more realistic models of the plasma membrane in Gram-negative and Gram-positive bacteria respectively, the conformation of the peptides in lipid bilayers, composed of either POPE/POPG (75:25 mol:mol) or POPG only, was investigated in the steady state through far-UV CD spectroscopy (Fig. S1). Small amounts of peptide stock solution were added to small unilamellar vesicles (SUVs) to decrease the lipid-to-peptide ratio and investigate the effect of peptide concentration on the secondary structure. Measurements were performed in the presence of 100 mM NaCl, to maintain as similar environment as possible to those used for other experimental procedures. All three peptides adopted a disordered structure when in buffered solution and adopted ordered conformations, of the type described for temporin B KKG6A in SDS. The titration indicates that all peptide molecules in solution bind to the vesicles. A notable exception in the series is illustrated for temporin B itself in the presence of POPE/POPG (75/25) and POPG, with a less pronounced negative 220 nm feature which indicates a more ordered tail or a less ordered β-turn (Fig. S1A/D). Otherwise, differences are small and the CD spectra can be said to be broadly similar.

### MD simulations reveal different conformational flexibilities on initial binding to model membranes

To characterise the interaction of temporin B analogues with model membranes with atomic resolution, duplicate all atom simulations were run for 100 ns. In each case, using the structures determined by NMR in SDS for the starting configuration, eight peptides for each temporin B analogue were placed in water above bilayers comprising either 512 POPG lipids or a mixture of 384 POPE and 128 POPG lipids.

Averaged psi dihedral angles for each residue in the eight temporin B peptides are similar in the presence of both POPE/POPG (Fig. 2A) and POPG (Fig. 2B) membranes; as in previous work,^11^ phi dihedral angles were not affected and are not shown. The values for these psi dihedral angles do not lie in the range of values accepted for α-helix conformations but instead correspond to more extended conformations.^41^ Temporin B displays considerable conformational flexibility during these first 100 ns which increases from N- to C-terminus. Modifications made to generate temporin B L1FK have a noticeable impact on the preferred conformation adopted in both membrane types; the deletion of Asn7, such that Leu7 now follows Gly6, induces a shift in local conformation with psi angles for residues 6-8 (GLL) consistent with the β-turn (single turn α-helix) conformation (Fig. 2C/D). The conformational flexibility of temporin B is however retained throughout the length of the modified peptide.

**Figure 2.**
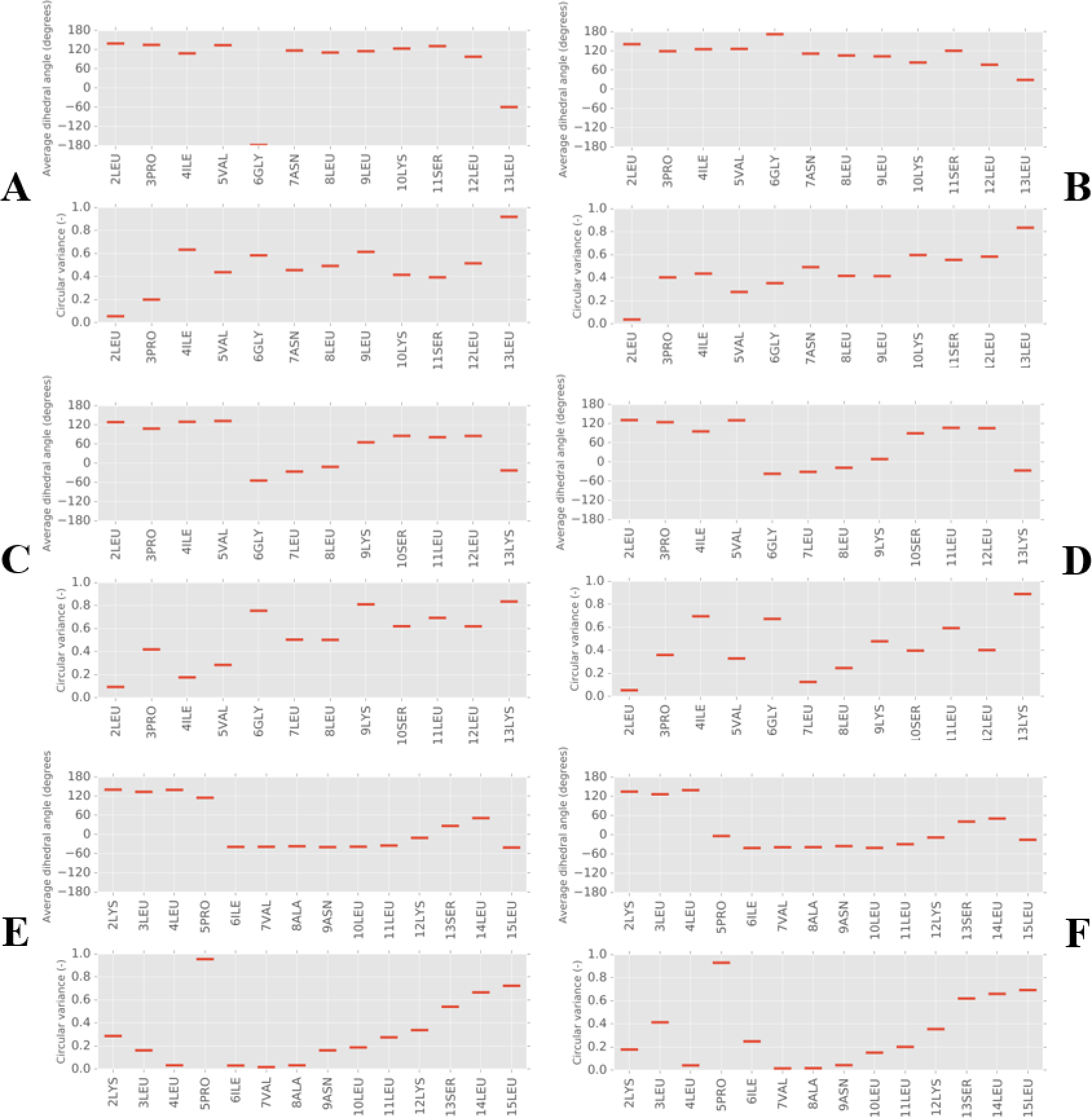
Secondary structure analysis of temporin B peptides from MD simulations in model membranes. Dihedral angles (psi) and their circular variance are shown for each residue averaged over 100 ns of simulation and eight peptides in temporin B (A/B), temporin B L1FK (C/D) and temporin B KKG6A (E/F) peptides when binding to POPE/POPG (A/C/E) or POPG (B/D/F) membranes.

The modifications made to generate temporin B KKG6A have a much more substantial impact. Except for the five residues at the N-terminus, there is a shift in psi angles associated with the replacement of a glycine (Gly6 in temporin B) with alanine (Ala9 in temporin B KKG6A) such that residues 6-12 (PIVANLK) adopt a two turn α-helix conformation (Fig. 2E/F). Conformational flexibility is also reduced throughout the length of the peptide with only the two residues at the C-terminus unaffected (Fig. 2E/F).

### Modification of temporin B affects its penetration of lipid bilayers and aggregation at their surface

In addition to changes in preferred secondary structure, MD simulations allow visualisation of the ability of the three temporin B peptides to penetrate either POPE/POPG or POPG bilayers (Fig. 3). All peptides start the simulation in water with no contact with the membrane. After the simulation had run for 100 ns all the peptides were interacting with the lipids regardless of the composition of the membrane. In general, the temporin B peptides showed deeper insertion in POPG membranes, as observed previously,^42^ compared with POPE/POPG membranes, with contact and insertion beginning at the N-terminus in all cases. The temporin B peptides were also frequently self-associated (Fig. 4) and assembled as dimers, trimers or even tetramers (Fig. S2-6). Aggregation was consistently lower in POPG membranes compared with POPE/POPG membranes.

**Figure 3.**
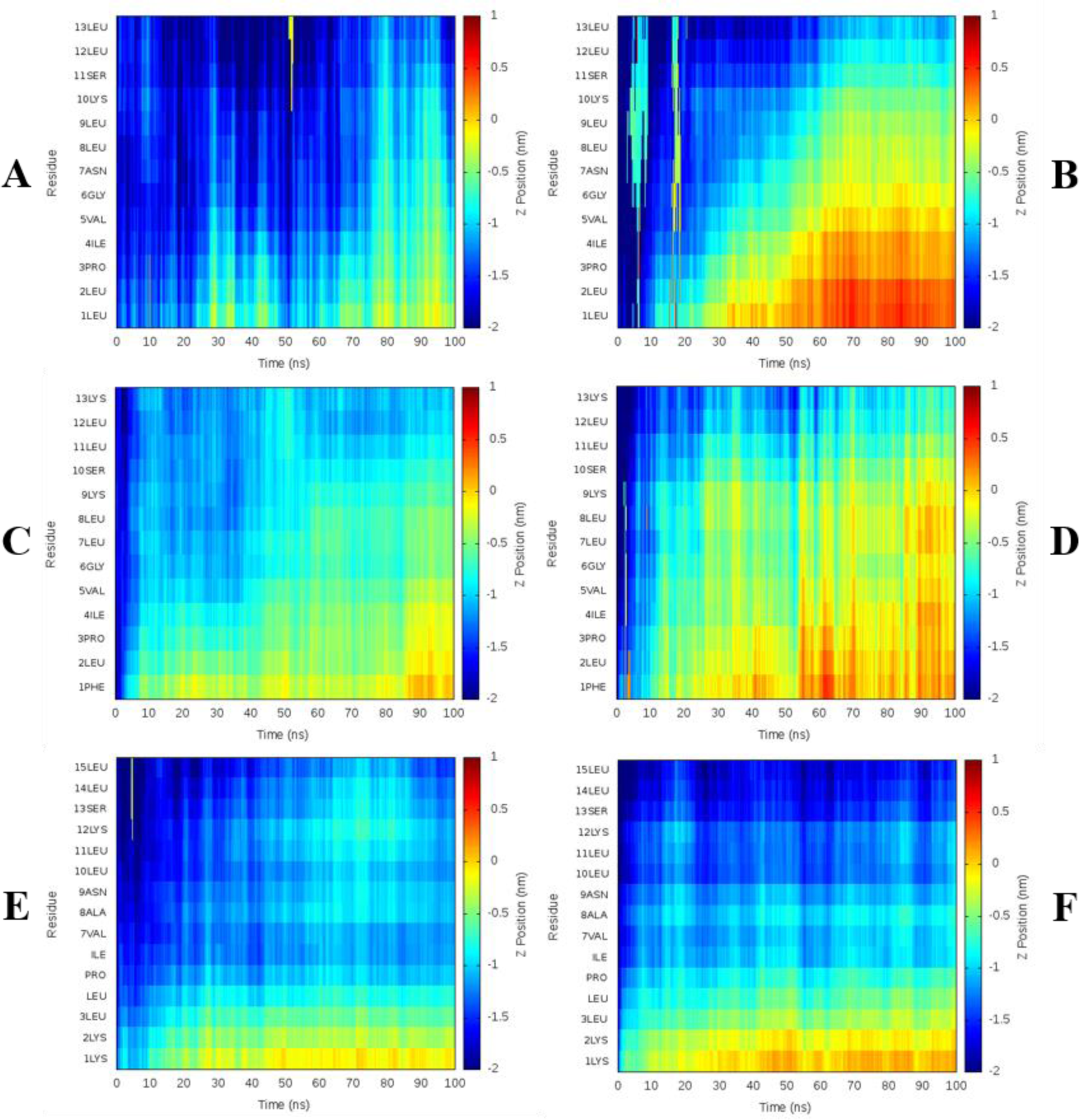
Temporin B peptide topology during 100 ns MD simulation in model membranes. The depth of temporin B (A/B), temporin B L1FK (C/D) and temporin B KKG6A (E/F) insertion is shown as the Z-position relative to the phosphate group for each residue averaged over all eight peptides in each simulation. Positive or negative values indicate the peptides are below or above the phosphate group, respectively for peptides in POPE/POPG (A/C/E) or POPG (B/D/F).

**Figure 4.**
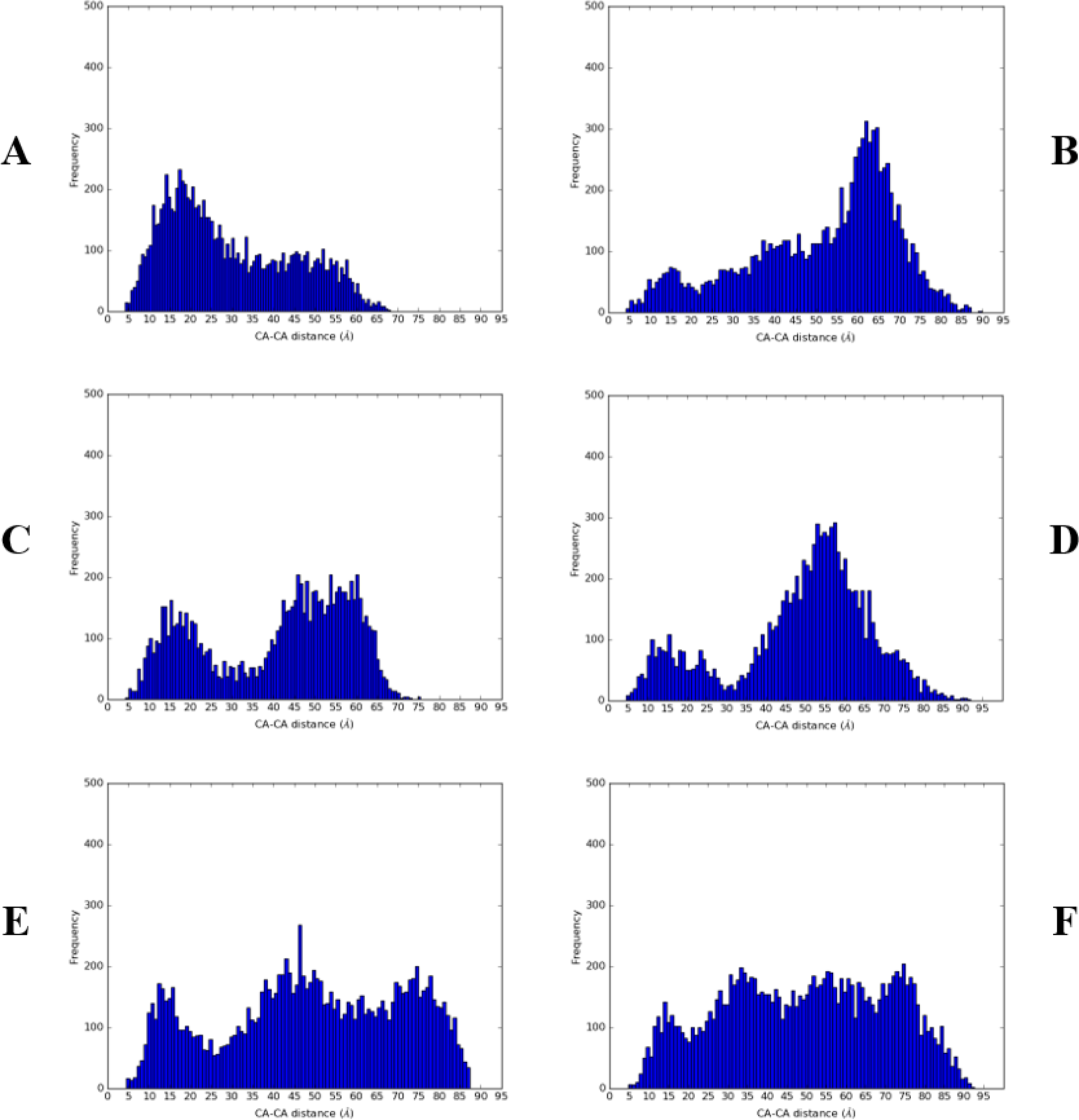
Temporin B peptides aggregation during 100 ns MD simulation in model membranes. The probability of temporin B (A/B), temporin B L1FK (C/D) and temporin B KKG6A (E/F) aggregation is reported as frequency histogram of the Cα-Cα distances, respectively for peptides in POPE/POPG (A/C/E) or POPG (B/D/F) membranes.

Temporin B is known to act in synergy with temporin L with the latter acting to inhibit the aggregation of the former.^43^ Acting alone, temporin B struggles to penetrate far into POPE/POPG membranes, with residues 1-7 only inserting and these reaching only the interfacial region of the bilayer (Fig. 3A), with most peptides residing in an aggregated state (Fig. 4A; Fig. S2). In POPG membranes the aggregation of temporin B is much less pronounced (Fig. 4B) and penetration proceeds more rapidly, is deeper – beginning to enter the hydrophobic core - and more of each peptide becomes buried (Fig. 3B; Fig. S3).

Temporin B L1FK behaves similarly to temporin B (Fig. 3CD; Fig. 4C/D). However, its initial binding and the insertion were more effective than observed for the parent peptide. Temporin B L1FK inserted almost flat in both lipid compositions enabling more of the peptide to be buried (Fig. 3C/D) and, notably in POPE/POPG, had a much lower tendency to self-associate than temporin B (Fig. 4A/C). Again, in POPG bilayers the tendency to self-associate is reduced (Fig. 4D) and penetration into the hydrophobic interior of the membrane is greater than in POPE/POPG bilayers (Fig. 3D; Fig S4/5).

In contrast with temporin B and temporin B L1FK, temporin B KKG6A fails to insert into either bilayer beyond the first two residues of the N-terminus (Fig. 3E/F). The result of additional positive charges at the N-terminus, the greater proportion of α-helix conformation and the reduced conformational disorder is that temporin B KKG6A has a much lower tendency to form higher order aggregates in either membrane, though may still form dimers and trimers (Fig. S6/7). Penetration mediated by the previously hydrophobic N-terminus is halted by interactions between the two lysines and anionic lipid headgroups.

### Peptide effects on membrane order

The relative differences in the ability of the temporin B peptides to penetrate lipid bilayers is expected to impact on their ability to disorder, disrupt or permeate membranes. In the MD simulations described above, temporin B and temporin B KKG6A had a greater tendency to cluster POPG lipids in mixed membranes (Fig. 5A).

**Figure 5.**
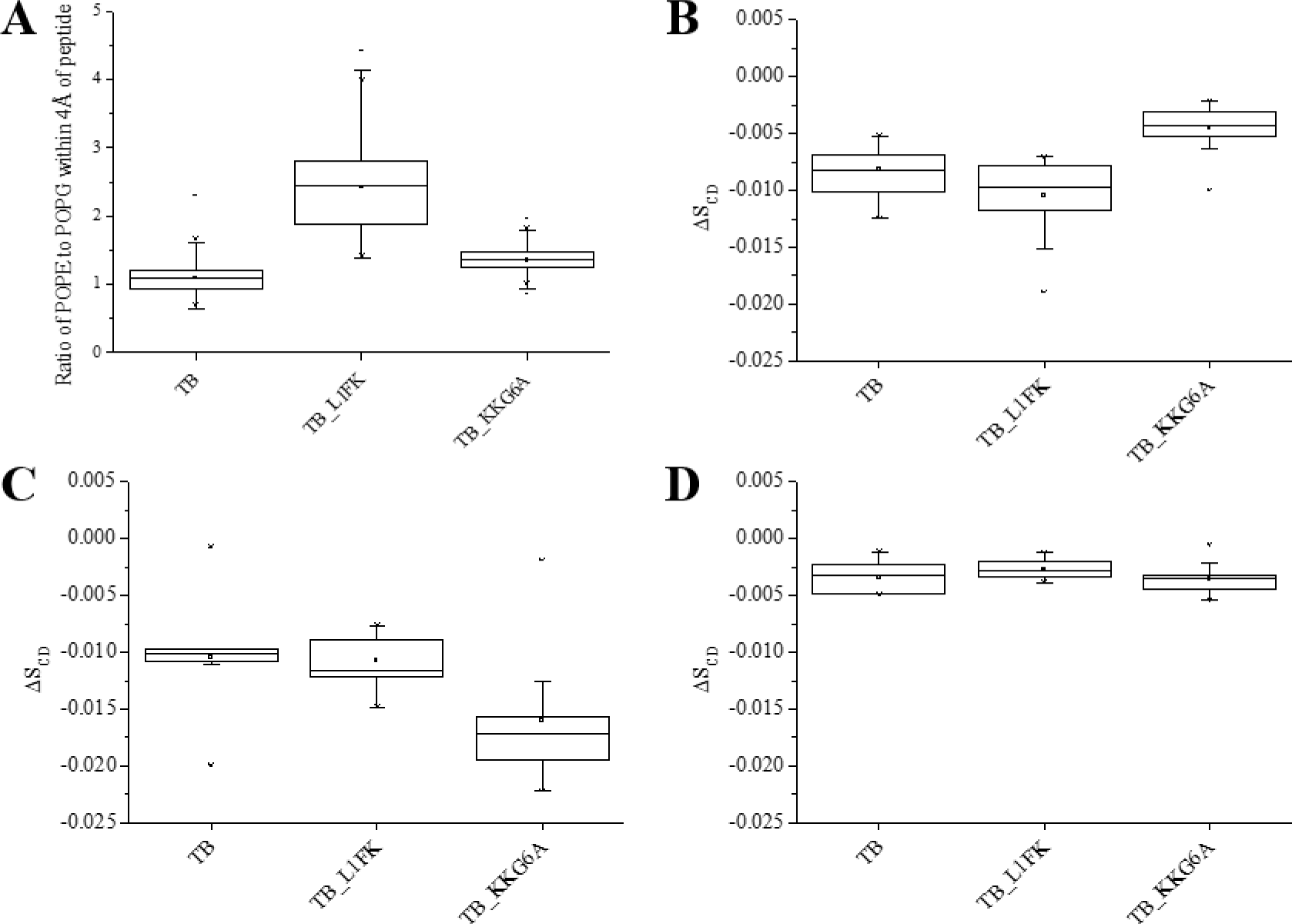
The effect of temporin B peptides on lipids. Capability of the peptides to cluster anionic lipids in POPE/POPG (75:25 mol:mol) membranes in silico (A). The POPE/POPG ratio of the lipids within 4Å from the peptides is compared across the three peptides where a ratio lower than three indicates the peptide tends to cluster the anionic lipids. Temporin B and temporin B KKG6A show a higher clustering effect. AMP disordering effect on lipid membranes in the steady state, as obtained from ^2^H solid-state NMR experiments (B-D). The effect of 2 mol% of each AMP was studied in multilamellar vesicles (MLVs) composed of POPE/POPG-d_31_ (75:25) (B), POPE/POPE-d_31_/POPG (50:25:25) (C) and POPG/POPG-d_31_ (75:25) (E) in 5 mM Tris-amine buffer, pH 7.00 and 100 mM NaCl.

This could be related either to the greater nominal positive charge of these peptides relative to temporin B (Table 1) or to their inability to penetrate easily beyond the interfacial region such that cationic residues remain concentrated at the bilayer surface and induce lateral separation of anionic and zwitterionic lipids. The interaction of cationic AMPs with anionic lipids has been demonstrated previously, *in vitro*^11,44,45^ and *in silico*^11,46,47^ to induce disordering of the lipid acyl chain and is associated with antimicrobial activity. Here, a good qualitative comparison was observed between the peptide induced membrane disordering obtained experimentally, in the steady-state, using ^2^H solid state NMR (Fig. 5B-D) and from disordering of lipids determined to be within a 4 Å ring of each peptide in MD simulations (Fig. S8).

In the steady state, the average difference (ΔS_CD_) between the order parameter in the presence and absence of 2 mol% of each temporin B peptide on the POPG-d_31_ (Fig. 5B) and POPE-d_31_ (Fig. 5C) component in POPE/POPG membrane and POPG-d_31_ in POPG membranes (Fig. 5D) allows comparison of any differing ability to disorder different components of each membrane. Notably, temporin B L1FK has a significantly (p < 0.05) greater disordering effect on the anionic component in mixed membranes than temporin B KKG6A (Fig. 5B) while the latter has a greater disordering effect on the zwitterionic POPE when compared with both temporin B and temporin B L1FK (p < 0.05) (Fig. 5.C).

No effect of any of the three peptides can be detected on the order parameter for POPG-d31 in POPG membranes (Fig. 5D) though the measured order parameter here is more likely to be reporting on an average of those lipids interacting with the peptides and those located in the remaining bulk than in the analogous experiments on mixed POPE/POPG-d31 membranes where a higher probability of the labelled anionic lipid associating with cationic AMP is expected.

### Temporin B analogues have drastically different abilities to induce conductance activity in model membranes

The whole-cell patch-clamp technique allows the measurements of current transitions on a phospholipid membrane. It has been largely used to study conductance and kinetics of membrane channels in cells and in lipid bilayers.^48^ The recent development of the Port-a-Patch^®^ automated patch-clamp system from Nanion Technologies (Munich, Germany), enables rapid testing of membrane active molecules with an automated system, bringing this technique into the drug discovery pipeline.^49–51^ Previous works have applied this technique to investigate the mode of action of AMPs.^52–59^ In particular, the three main membrane-activity models described in the introduction can be distinguished.^52^ This protocol involved the use of a microperfusion system on a standard pipette patch-clamp set up and mammalian cells.^52^ We report here that the same aim was achieved with the Port-a-Patch® system and the use of model membranes, which enabled the peptides to be distinguished on the basis of their disruption of either DPhPE:DPhPG (60:40 mol:mol) or DPhPG bilayers, mimicking Gram-negative and Gram-positive bacteria, respectively.^60–64^ Diphytanyl chains are used here since membranes composed of these lipids are in a fluid phase at room temperature, irrespective of headgroup.

The membrane activity of the peptides can be distinguished on the basis of 1) the concentration required for membrane activity to be recorded (Table 2), 2) the latency – the time taken for conductance to begin after addition of peptide, 3) the duration of conductance activity, 4) the amplitude and number of each conductance event and 5) whether the AMP ultimately caused the membrane to break.

**Table 2.** Concentration of peptide necessary to start membrane activity in electrophysiology experiments.

| Peptide | Peptide concentration (μM) |  |
| --- | --- | --- |
|  | DPhPE/DPhPG | DPhPG |
| Temporin B | 25.34 | 35 |
| Temporin B L1FK | 11.32 | 4.94 |
| Temporin B KKG6A | 200 | 100 |

The concentration used for the experiments for each peptide is the minimum concentration necessary to observe activity and was obtained with a peptide titration of the model membrane (Table 2). Notably there is an apparent relationship between the ability of each temporin B peptide to insert rapidly into model membranes, as determined by the MD simulations, and the concentration required to induce conductance. Less than half as much temporin B L1FK as temporin B was required to induce conductance in DPhPE/DPhPG membranes while approximately seven times more temporin B than temporin B L1FK is required to induce conductance in DPhPG membranes. Temporin B KKG6A, for which no more than the two N-terminal residues penetrated either POPE/POPG or POPG bilayers, required nearly eighteen and over twenty times as much peptide as temporin B L1FK to induce conductance in, respectively, DPhPE/DPhPG and DPhPG membranes.

Characteristic channel like activity - well-defined events with discrete opening levels – has been observed previously by others for alamethicin, gramicidin A, cecropin B and dermicidin^53–55^ and for other peptides in our hands. However, Temporin B presented an irregular activity in both mixed and negatively charged membranes (Fig. 6A/B). Temporin B caused a perturbation of the membrane characterized by bursts of current (fast events seen mostly as spikes) with a wide distribution of amplitudes, ultimately resulting in a destabilization of the membrane. Interestingly, temporin B activity started relatively soon after peptide addition, always causing an immediate rupture of the membrane (Fig. 7).

**Figure 6.**
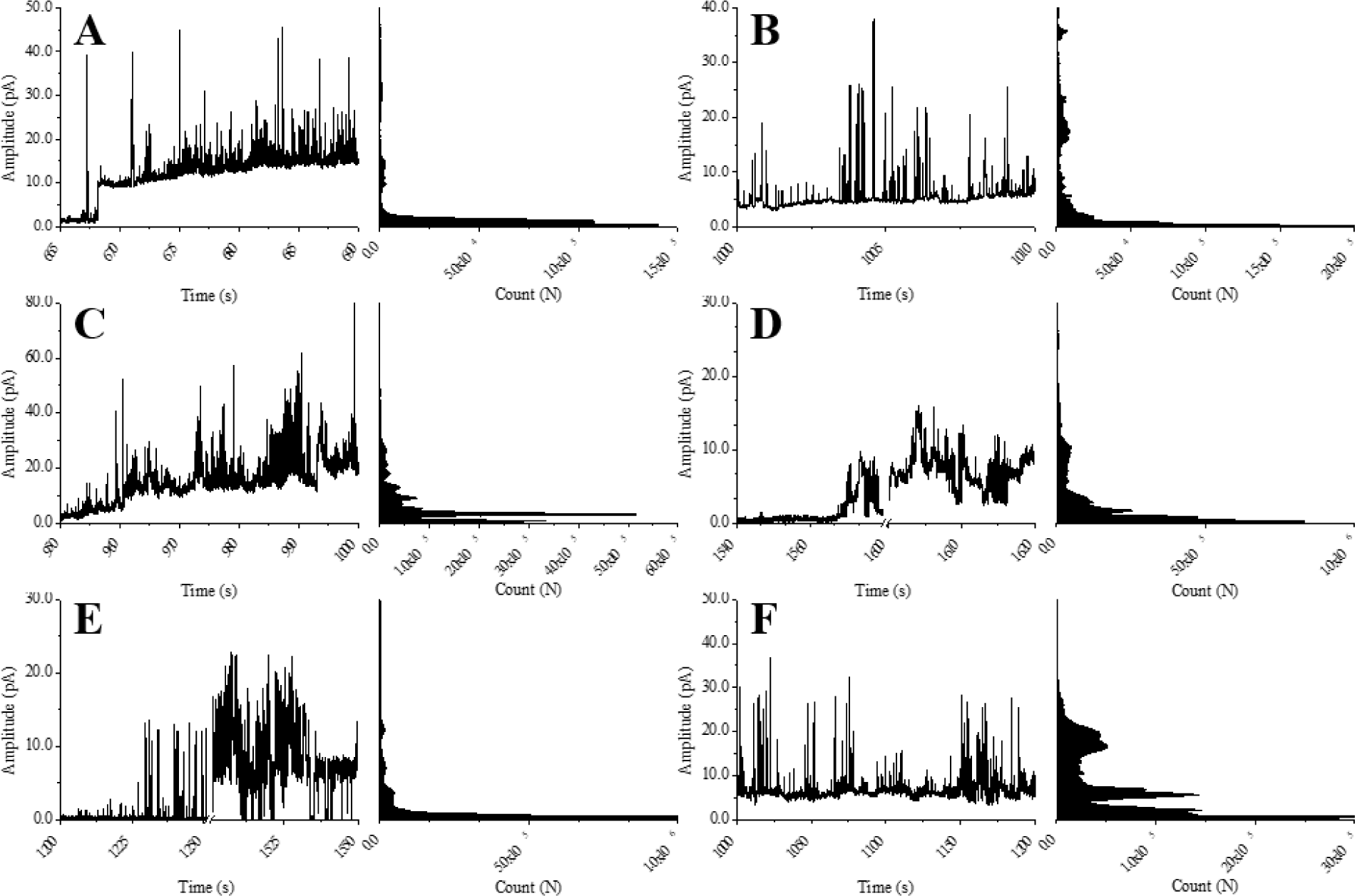
Membrane activity of the temporin B analogues. Representative current traces and all-points histograms for temporin B (A/B), temporin B L1FK (C/D) and temporin B KKG6A (E/F) in DPhPE/DPhPG (A/C/E) or DPhPG (B/D/F) membranes. Experiments were acquired with a holding potential of +50 mV and in the presence of 250 mM KCl, 50 mM MgCl_2_, 10 mM HEPES, pH 7.0. Y-axis is scaled accordingly to the different amplitudes for the different peptides.

**Figure 7.**
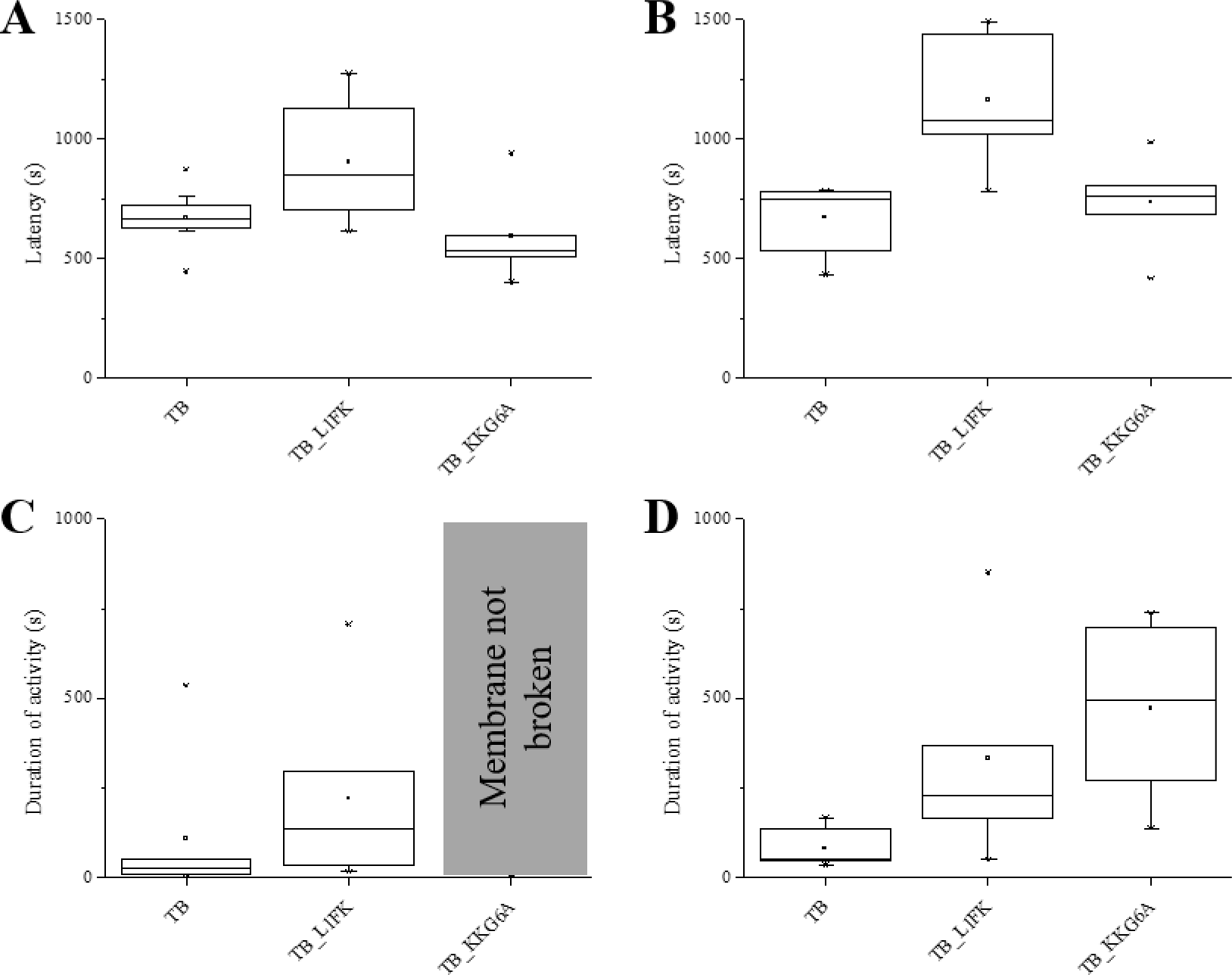
Latency and duration of membrane activity. The average time elapsed between peptide addition and the first appearance of membrane activity (A/B) and the average duration of activity before the membrane breaks (C/D) is shown for DPhPE/DPhPG (A/C) and DPhPG (B/D) membranes.

Conductance induced by temporin B L1FK was also unaffected by membrane composition. Temporin B L1FK acted more slowly than temporin B (significantly so in DPhPG; p < 0.05) and induced a mixture of membrane activities, characterised by some channel-like events, with defined levels of amplitude, and irregular, fast events, similar to those induced by temporin B (Fig. 6C). In DPhPG, more pore-like events were visible, however the distribution of the opening and closing levels was broad and no single populations could be defined (Fig. 6D). Temporin B L1FK often broke DPhPE/DPhPG membranes, while it rarely broke DPhPG membranes (Fig. 7C/D), perhaps since conductance was achieved with a lower active concentration.

Membrane conductance induced by temporin B KKG6A was not detectable by electrophysiology unless a high concentration was reached. At these high concentrations, considerable destabilization of the membrane is observed with more numerous high amplitude events than observed for temporin B, particularly in DPhPG (Fig. 6E/F). Although conductance appears relatively soon after temporin B KKG6A addition, the membrane remains resilient and either does not break (Fig. 7C) or takes significantly longer (p < 0.05) to break than when challenged with much less temporin B (Fig.7D).

## Discussion

### Time-resolved techniques distinguish mechanistic features between the three temporin B peptides

Here we investigated whether a combined biophysical approach is capable of discriminating the species specific antibacterial mechanisms of action between three analogues of temporin B. Although structures for the three peptides solved in anionic SDS were noticeably different, when applied to model membranes, little difference in conformation could be discerned in the steady state. To address this, we developed an approach using the Port-a-Patch® tool from Nanion Technologies (Munich, Germany) and correlated the findings obtained from patch-clamp analysis to the primary sequence and the biophysical behaviour of the peptides. Notably, these two time-resolved techniques were better able to distinguish the activity of the temporin B analogues. The time course of AMP action is increasingly recognized as an important parameter and it is notable that the time taken for membrane conductance to appear in model membranes is consistent with the time taken for the attack of AMPs on bacteria reported elsewhere.^65,66^ Although the pharmacodynamics of differing peptides will be variable, the latency associated with membrane activity of between 10 and 20 minutes for temporin B analogues is consistent with permeabilization of the *E. coli* cytoplasmic membrane^65^ and the induction of oxidative stress^66^ by LL-37 after around 25 minutes, well after bacterial growth has halted. It is not possible to sample such timescales using all-atom molecular dynamics simulations. Instead, the MD simulations here describe the initial binding of the temporin B analogues to model bilayers and succeed in discriminating different binding and insertion which can be directly related to the differences in primary sequence.

The study of membrane active molecules, such as AMPs, with electrophysiology is complicated by their irregular activity when interacting with the bilayers. Indeed, only few AMPs apparently form regular single-channels when analysed with this technique,^53–55^ while many show irregular membrane activity which is hard to define.^53,67^ Nevertheless, a rationale has been suggested in order to distinguish between the “barrel-stave”, “toroidal” pores or “carpet-like” mechanisms on the base of the current traces.^52^ Events showing a single-channel nature would characterise a pore, while an irregular current trace would be representative of a carpet mechanism. Moreover the different mechanisms could be distinguished on the basis of shorter or longer time constants of current activation and recovery, indicating a “barrel-stave” or a “toroidal” pore, respectively, and the irreversibility of the permeabilization process that would be the signature of a “carpet” like mechanism (detergent-like).^52,56^ This characterisation is based on the predominant models currently used to describe AMPs activity.^1,7^ However, more recent studies indicate many events at the bacterial plasma membranes are poorly described by these existing models and refinements or entirely new models are required to satisfactorily explain the behaviour of some AMPs; for instance the “snorkelling mechanism”,^68,69^ the “dynamic peptide-lipid supramolecular pore”^67,70–72^ or the “disordered toroidal pore”.^42,47,73^ Informed by molecular level information provided by MD simulations, the substantial changes in membrane activity amongst the three temporin B analogues are discussed below and indicate that relatively minor modifications of the primary sequence can cause substantial shifts between behaviours associated with each of the models described above.

### Temporin B is a hydrophobic AMP, prone to self-association

Temporin B and its two analogues are poorly studied, and little information is known about their mode of action. Hydrophobic AMPs such as temporin B have been proposed to go through an aggregation process and form fibrils as a step in the bactericidal process.^74^ As with amyloid peptides, this mechanism would form a cytotoxic intermediate which would cause the death of the cell (bacterial in this case) through poration of the plasma membrane. Evidence is indeed provided here for substantial aggregation of temporin B and its analogues but there is no evidence of any conformational shift, from α-helix to β-sheet, as characteristic of amyloid-like peptides as was predicted.^74^ A self-association of temporin B monomers through hydrophobic interactions was also reported on lipopolysaccharide;^37,75^ this was suggested to be the cause of the poor activity of this peptide against Gram-negative bacteria. Further, an increase in activity was observed for temporin B when used in combination with temporin L with the latter reported to inhibit the self-association. Presumably, temporin L can stabilise the monomer conformation of temporin B and/or cause structural changes in lipopolysaccharide when bound, which prevents temporin B aggregation. When in combination, the two peptides showed a higher potency in lipopolysaccharide disintegration.^13,37,75^ Primary sequence modification strategies that seek to reduce self-association of temporin B may therefore enhance its activity against Gram-negative bacteria.

### Modification of the temporin B primary sequence to create temporin B L1FK

Temporin B L1FK was computationally designed with a statistical model considering the cytotoxicity of the sequence, but also a close physicochemical pattern to the parent peptide.^16^ Temporin B L1FK behaves similarly to temporin B in inducing conductance albeit with a greater number of channel-like events and at somewhat lower concentrations and this correlates with the improved potency obtained against both *E. coli* NCTC 9001 and *S. aureus* Newman. We ascribe this improvement in membrane activity, and consequently in anti-bacterial potency, to increased insertion into the membrane associated with reduced self-association. This is achieved with greater hydrophobicity at the N-terminus by substituting phenylalanine for leucine, the increased positive charge and the reduced conformational flexibility between residues six and eight. For both temporin B and temporin B L1FK, patch-clamp revealed no difference in activity according to membrane composition. In neither case could the observed activity be defined as regular single-channel formation, but rather a perturbation of the permeability of the membrane due to defects of the lipid bilayer that cause spikes of current. Unlike temporin B, temporin B L1FK, in addition to the spikes also caused some square-shaped events resembling channels or pores but characterized by disordered openings in which an uneven distribution of levels can be detected. Such conductance activity is well removed from that reported for “barrel-stave” or “toroidal” pores, as described previously for alamethicin, gramicidin A, cecropin B and dermicidin.^53–55^ Instead, the current traces obtained for both temporin B and temporin B L1FK are comparable to the ones obtained for gramicidin S,^53^ for which they suggest a mechanism similar to the so called “in-plane diffusion” model.^76,77^ In this model the peptides insert into the phospholipid bilayer disordering the packing and causing defects that can increase the membrane permeability. However, in this model the aggregation of peptides is considered minimal and unfavourable, while we observed considerable self-association for temporin B and, although reduced, also temporin B L1FK. The immediate membrane rupture observed for temporin B in all the electrophysiology measurements is consistent with the “carpet-like” mechanism, where peptides disintegrate the membrane following a detergent-like process.^56,78^ However, such behaviour was not observed for temporin B L1FK. Current traces similar to those obtained for temporin B L1FK, characterised by abrupt transient current spikes, were reported for poly-arginine peptides, known to act as cell-penetrating peptides (CPP).^79^ The deep and flat insertion, observed for this temporin B during the MD simulations, might suggest the ability of this peptide to better penetrate the bilayer and act inside the cell – activity not captured by techniques applied in the present study.

### Modification of the temporin B primary sequence to create temporin B KKG6A

Modification of temporin B through addition of two lysines at the N-terminus and replacement of a glycine with an alanine ensures that self-association of the resulting peptide, temporin B KKG6A, is substantially reduced. This is achieved though at cost of preventing insertion of the peptide, beyond the first two residues at the N-terminus, into either membrane and a very large increase in the amount of peptide required to obtain any conductance activity. This is a reminder that not only the net charge, but also its distribution along the sequence is important for AMPs activity.^19,73^ However, while this might explain the loss of activity against *S. aureus*, activity against *E. coli* was actually improved compared to the original temporin B peptide. Notably however, and in contrast to temporin B and temporin B L1FK that disorder the anionic phosphatidylglycerol component, temporin B KKG6A disorders the zwitterionic phosphatidylethanolamine component of membranes designed to mimic the plasma membrane of Gram-negative bacteria. This, together with a clear interaction with the headgroup region of the phospholipid bilayer observed in the MD simulations, suggests that the peptide is interacting with the membrane, but that the effect of this interaction does not readily induce channel conductance detectable through patch-clamp. The disorder observed in phosphatidylethanolamine lipids is related to the observation, in MD simulations, that temporin B KKG6A is particularly effective at clustering anionic lipids. This can be toxic to bacteria by disrupting membrane domains essential for specific interactions with other molecules^80^ and might not cause a leakage visible with technique as patch-clamp. Adoption of this mechanism against *E. coli* is consistent with the loss of activity against Gram-positive *S. aureus*, the plasma membrane of which is mainly composed of negatively charged lipids.^60–64^ Whether this or other mechanisms more accurately describe the behaviour and anti-bacterial activity of temporin B KKG6A, it is clear that, following the optimization process, its mechanism of action differs substantially from the native temporin B peptide.

## Conclusion

Since cecropin, the first AMP, was isolated from the hemolymph of the silk moth Hyalophora cecropia in 1981,^81^ AMPs have been studied as a new interesting class of molecules for the development of new antibiotics. The attraction of these short, cationic peptides is related to their ability to defeat infections from many different pathogens (bacteria, fungi, viruses) and, in particular, the low rate of bacterial resistance onset.^1–3,6^ Despite the efforts made to characterise different AMPs from different sources, there is still a lack of understanding of their specific modes of action and variation thereof due to sequence modifications.^1^ The activity of temporin B is likely improved *in vivo* through its synergistic interaction with temporin L which acts to prevent self-association. Both sequence modification strategies described here also act to reduce self-association of the AMP in the target membrane but lead to fundamentally different behaviour which was revealed by a combination of MD simulations and patch-clamp conductance recordings but not by e.g. circular dichroism. Therefore, in characterising the membrane interaction of temporin B and its analogues, the present study shows that even minor alterations to AMP primary sequences may cause drastic changes in the mechanism of interaction with bacterial plasma membranes which in turn affect antibacterial potency and selectivity. Future attempts to enhance the potency of designed or naturally occurring AMP sequences will be aided by as sophisticated as possible understanding of the crucial peptide-peptide and peptide-membrane interactions and their outcomes.

## ASSOCIATED CONTENT

### Supporting Information

CD spectra of temporin B peptides in model membranes obtained in the steady-state, snapshots and analysis of aggregation state and temporin B peptide induced lipid disordering during MD simulations are available together as Supporting Information.

## AUTHOR INFORMATION

### Author Contributions

GM, CDL, DAP and AJM designed the study. GB provided materials. GM, PMF and AJM wrote the main manuscript text and prepared all figures. PMF and CDL performed and/or analysed atomistic simulation data. GM and RAA performed NMR structural studies. GM, VBG and AJM performed solid-state NMR. GM, VBG, TTB and AFD performed and/or analysed CD measurements. GM and VBG performed and analysed patch-clamp measurements. All authors approved the manuscript.

### Notes

The authors declare no competing interests.

## Acknowledgement

NMR experiments described in this paper were produced using the facilities of the Centre for Biomolecular Spectroscopy, King’s College London, acquired with a Multi-user Equipment Grant from the Wellcome Trust and an Infrastructure Grant from the British Heart Foundation. CDL. acknowledges the stimulating research environment provided by the EPSRC Centre for Doctoral Training in Cross-Disciplinary Approaches to Non-Equilibrium Systems (CANES, EP/L015854/1). PMF is supported by a Health Schools Studentship funded by the EPSRC.

